# Sex Differences In The Interaction Between Alcohol And mTORC1

**DOI:** 10.1101/2023.10.04.560781

**Authors:** Yann Ehinger, Khanhky Phamluong, Dorit Ron

**Author notes:** Correspondence: D. Ron, Department of Neurology, University of California, San Francisco, 675 Nelson Rising Lane, BOX 0663, San Francisco, CA 94143-0663, USA.

## Abstract

The kinase mechanistic target of rapamycin complex 1 (mTORC1) plays an essential role in learning and memory by promoting mRNA to protein translation of a subset of synaptic proteins at dendrites. We generated a large body of data in male rodents indicating that mTORC1 is critically involved in mechanisms that promote numerous adverse behaviors associated with alcohol use disorder (AUD) including heavy alcohol use. For example, we found that mTORC1 is activated in the nucleus accumbens (NAc) and orbitofrontal cortex (OFC) of male mice and rats that were subjected to 7 weeks of intermittent access to 20% alcohol two-bottle choice (IA20%2BC). We further showed that systemic or intra-NAc administration of the selective mTORC1 inhibitor, rapamycin, decreases alcohol seeking and drinking, whereas intra-OFC administration of rapamycin reduces alcohol seeking and habit in male rats. This study aimed to assess mTORC1 activation in these corticostriatal regions of female mice and to determine whether the selective mTORC1 inhibitor, rapamycin, can be used to reduce heavy alcohol use in female mice. We found that mTORC1 is not activated by 7 weeks of intermittent 20% alcohol binge drinking and withdrawal in the NAc and OFC. Like in males, mTORC1 signaling was not activated by chronic alcohol intake and withdrawal in the medial prefrontal cortex (mPFC) of female mice. Interestingly, Pearson correlation comparisons revealed that the basal level of mTORC1 activation between the two prefrontal regions, OFC and mPFC were correlated and that the drinking profile predicts the level of mTORC1 activation in the mPFC after 4-hour binge drinking. Finally, we report that administration of rapamycin does not attenuate heavy alcohol drinking in female animals. Together, our results suggest a sex-dependent contribution of mTORC1 to the neuroadaptation that drives alcohol use and abuse.

## INTRODUCTION

The kinase mechanistic target of rapamycin (mTOR) is a serine and threonine kinase that when localized in a complex that includes Raptor, Deptor and mLST8, is termed mTOR Complex 1 (mTORC1) (Lipton and Sahin 2014, Saxton and Sabatini 2017). mTORC1 is activated by stimuli such as growth factors, amino acids, and oxygen (Sengupta, Peterson et al. 2010). mTORC1 which is activated mainly through the H-Ras/Phosphatidylinositol 3-Kinase (PI3K)/AKT pathway (Hay and Sonenberg 2004), plays an important role in lipid genesis, glucose homeostasis, protein translation and autophagy (Kim and Guan 2015, Saxton and Sabatini 2017, Liu and Sabatini 2020). In the mature brain, mTORC1 is activated by neurotransmitters and neuromodulators, such as glutamate and BDNF (Lipton and Sahin 2014, Morisot, Phamluong et al. 2019, Conde-Dusman, Dey et al. 2021). Once activated, mTORC1 phosphorylates eIF4E-binding protein (4E-BP) and the ribosomal protein S6 kinase (S6K), which in turn phosphorylates its substrate, S6 (Saxton and Sabatini 2017). mTORC1, 4E-EBP, S6K and S6 are part of the ribosomal translation machinery found in the CNS in both the cell bodies and dendrites (Buffington, Huang et al. 2014). The activation of the ribosomal translational machinery leads to the initiation of the translation of a subset of mRNA to proteins (Buffington, Huang et al. 2014, Santini, Huynh et al. 2014). mTORC1 plays an important role in synaptic plasticity, and learning and memory (Costa-Mattioli and Monteggia 2013, Lipton and Sahin 2014) and disruption in the normal functions of mTORC1 is linked to aging (Johnson, Rabinovitch et al. 2013), neurodegenerative diseases (Bove, Martinez-Vicente et al. 2011, Costa-Mattioli and Monteggia 2013, Liu and Sabatini 2020), neurodevelopmental disorders such as autism as well as psychiatric disorders including addiction (Costa-Mattioli and Monteggia 2013, Johnson, Rabinovitch et al. 2013, Neasta, Barak et al. 2014). Using male rodents as a model system, we generated data to suggest that mTORC1 plays an important role in neuroadaptations that underlie numerous AUD phenotypes in male rodents. Specifically, we found that both the first drink of alcohol (Beckley, Laguesse et al. 2016) as well as 7 weeks of IA20%2BC (Neasta, Ben Hamida et al. 2010, Laguesse, Morisot et al. 2017, Ehinger, Zhang et al. 2021) activate mTORC1 in the NAc. We showed that the cellular consequence of alcohol-mediated activation of mTORC1 in the NAc is the translation of synaptic proteins including proteins that promote F-actin (Laguesse, Morisot et al. 2017) and microtubules formation (Liu, Laguesse et al. 2017). We further surveyed the activation profile of the kinase in corticostriatal brain regions in male mice and rats and reported that alcohol-mediated mTORC1 activation is localized to the NAc shell and the OFC (Laguesse, Morisot et al. 2017). Using the selective mTORC1 inhibitors, Rapamycin and RapaLink-1 (Dowling, Topisirovic et al. 2010, Rodrik-Outmezguine, Okaniwa et al. 2016), we showed that mTORC1 in male rodents plays an important role in mechanisms underlying alcohol seeking, excessive alcohol consumption (Neasta, Ben Hamida et al. 2010), (Beckley, Laguesse et al. 2016, Morisot, Novotny et al. 2018), alcohol habit (Morisot, Phamluong et al. 2019), and the reconsolidation and reinstatement of alcohol reward memories (Barak, Liu et al. 2013, Ben Hamida, Laguesse et al. 2019). More recently, we used a binary drug strategy in which male mice were co-administered the mTORC1 inhibitor RapaLink-1, together with a novel small molecule (RapaBlock) that protects mTORC1 activity in the periphery, and showed that the co-administration of Rapablock with RapaLink-1 effectively reduced heavy drinking and seeking while eliminating RapaLink-1-dependent side effects (Ehinger, Zhang et al. 2021), suggesting that this approach may be used in subjects suffering from AUD. Since prior studies were conducted in male rodents, we set out to monitor mTORC1 activation pattern in the NAc, the OFC as well as the mPFC of female mice that were subjected to repeated cycles of binge alcohol drinking of 20% alcohol and withdrawal. We also examined the utility of rapamycin to reduce alcohol binge drinking and preference in female mice.

## METHODS

### Materials

Phosphatase Inhibitor Cocktails 2 and 3, and Dimethyl sulfoxide (DMSO) were purchased from Sigma Aldrich (St. Louis, MO). Nitrocellulose membrane was purchased from EMD Millipore (Billerica, MA, USA). Enhanced Chemiluminescence (ECL) was purchased from GE Healthcare (Pittsburg, PA). EDTA-free complete mini Protease Inhibitor Cocktail was purchased from Roche (Indianapolis, IN). NuPAGE Bis-Tris precast gels and Phosphate buffered saline (PBS) were purchased from Life Technologies (Grand Island, NY). Bicinchoninic Acid (BCA) protein assay kit was obtained from Thermo Scientific (Rockford, IL). ProSignal Blotting Film was purchased from Genesee Scientific (El Cajon, CA). Ethyl alcohol (190 proof) was purchased from VWR (Radnor, PA). Rapamycin (R-5000) was purchased from LC Laboratories (Woburn, MA).

### Antibodies

|  |  |  |
| --- | --- | --- |
| Mouse IgM anti-Tubulin | Santa Cruz Biotechnology (Dallas, TX) | SC-8035 |
| Rabbit anti-phospho S6 ribosomal protein | Cell Signaling (Danver, MA) | S235/236 |
| Rabbit anti- S6 ribosomal protein | Cell Signaling (Danver, MA) | #2708S |
| Donkey anti-mouse horseradish peroxidase | Jackson ImmunoResearch (West Grove, PA) | 715-035-150 |
| Donkey anti-rabbit horseradish peroxidase | Jackson ImmunoResearch (West Grove, PA) | 711-035-152 |

### Animals

Female C57BL/6J mice (6-8 weeks old at time of purchase) were obtained from The Jackson Laboratory. Mice were individually housed on paper-chip bedding (Teklad #7084), under a reverse 12-hour light-dark cycle (lights on 1000 to 2200 h). Temperature and humidity were kept constant at 22 ± 2°C, and relative humidity was maintained at 50 ± 5%. Mice were allowed access to food (Teklad Global Diet #2918) and tap water *ad libitum*. All animal procedures were approved by the university’s Institutional Animal Care and Use Committee and were conducted in agreement with the Association for Assessment and Accreditation of Laboratory Animal Care.

### Western blot analysis

Animals were killed and the brain was rapidly removed on an anodized aluminum block on ice. The NAc was isolated from a 1 mm thick coronal section located between +1.7 mm and +0.7_Jmm anterior to bregma, the OFC (3.1 mm and 2.1 mm) and mPFC (2.1 mm and 1.1 mm) according to the Franklin and Paxinos stereotaxic atlas (3rd edition). Collected tissues were immediately homogenized in 300 µl radioimmuno precipitation assay buffer containing (in mM: 50 Tris-HCl, pH 7.6, 150 NaCl, 2 EDTA), and 1% NP-40, 0.1% SDS and 0.5% sodium deoxycholate and a cocktail of protease and phosphatase inhibitors. Samples were homogenized by a sonic dismembrator, and protein content was determined using a BCA kit. Equal amounts of homogenates from individual mice (30 µg) were resolved on NuPAGE Bis-Tris gels and transferred onto nitrocellulose membranes. Blots were blocked in 5% milk-PBS, 0.1% Tween 20 for 30 min and then incubated overnight at 4_J°C with primary antibodies. Membranes were then washed and incubated with HRP-conjugated secondary antibodies for 2 hours at room temperature. Bands were visualized using ECL. The optical density of the relevant band was quantified using ImageJ 1.44c software (NIH).

### Preparation of solutions

Alcohol solution was prepared from absolute anhydrous alcohol (190 proof) diluted to 20% alcohol (v/v) in tap water. Rapamycin was dissolved in 3% DMSO and given intraperitoneally (i.p.) at a dose of 20 mg/kg (Laguesse, Morisot et al. 2017). Vehicle contained 3% DMSO.

### Alcohol drinking paradigm

Female mice underwent 7 weeks of IA20%2BC as described previously (Ehinger, Zhang et al. 2021). Specifically, mice had 24-hour access to one bottle of 20% alcohol and one bottle of water on Mondays, Wednesdays, and Fridays, with alcohol drinking sessions starting 2 hours into the dark cycle. During the 24 or 48 hours (weekend) of alcohol withdrawal periods, mice had access to a bottle of water. The placement (right or left) of the bottles was alternated in each session to control for side preference. Two bottles containing water and alcohol in an empty cage were used to evaluate the spillage. Alcohol and water intake were measured at the 4- and 24-hour time points. Alcohol preference ratio was calculated as the volume of alcohol intake/total volume of fluid intake (water_J+_Jalcohol).

To determine the effect of Rapamycin on alcohol drinking, mice were systemic administered with Rapamycin (20mg/kg) or vehicle 3 hours before the beginning of the drinking session, and alcohol and water intake were measured at the 4- and 24-hour time points.

### Statistics

GraphPad Prism 9 (GraphPad Software Inc., La Jolla, CA) was used to plot and analyze the data. D’Agostino–Pearson normality test and F-test/Levene tests were used to verify the normal distribution of variables and the homogeneity of variance, respectively. Data were analyzed using the appropriate statistical test, including two-tailed paired t test and one-way ANOVA followed by post hoc tests as detailed in Figure Legends. Correlation matrix computing the Pearson’s correlation of each variable with each other variable and heatmap displaying the Pearson correlation coefficient were performed in GraphPad Prism 9. All data are expressed as mean ± SEM, and statistical significance was set at p _J<_J 0.05.

## RESULTS

First, we examined whether alcohol activates mTORC1 signaling in the NAc of female mice. To do so female mice underwent 7 weeks of IA20%2BC as described in (Ehinger, Zhang et al. 2021) during which animals consumed large quantities of alcohol (4.5 ± 0.74 g/kg/4 hours, 17.21 ± 1.17 g/kg/24 hours (**Table 1**, **Fig. 1a**). On week 8, the NAc was dissected at the end of the 4-hour binge drinking session or at the end of the last 24-hour alcohol withdrawal session (**Fig. 1b**), and mTORC1 activity in alcohol or water only drinking female mice was assessed by determining the level of S6 phosphorylation which was used as a marker of mTORC1 activity.

**Figure 1:**
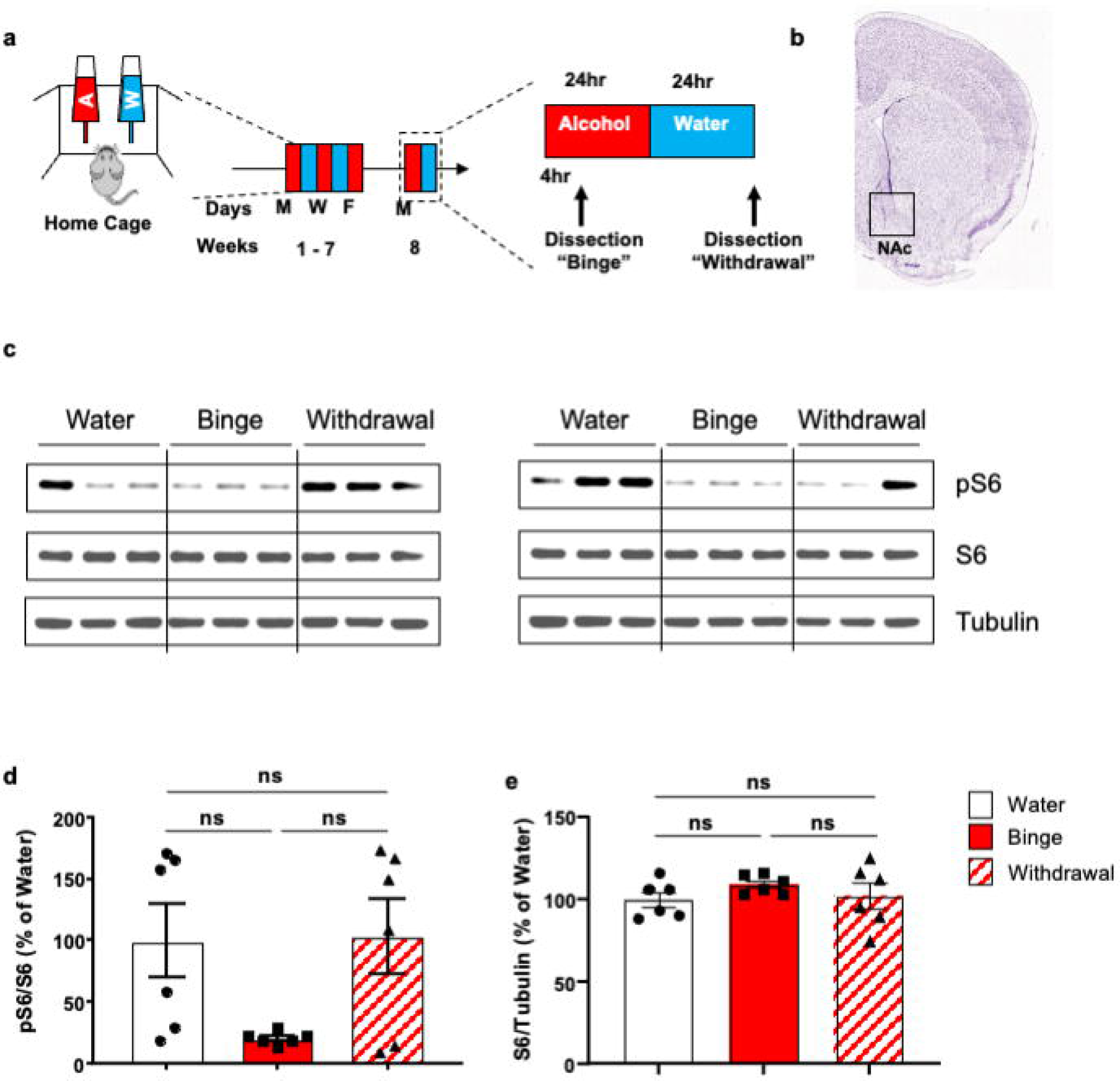
mTORC1 is not activated in the NAc of female mice in response to excessive alcohol intake. (**a**) Timeline of experiment. Mice underwent 7 weeks of IA20%-2BC whereas control animals had access to water bottles only. The nucleus accumbens (NAc) (**b**) was dissected either after the last 4h binge drinking session or at the end of the last withdrawal period and was subjected to western blot analysis. (**c-e**) PhosphoS6 and S6 protein levels were determined by western blot analysis. Tubulin was used as a loading control. ImageJ was used for optical density quantification. Data are presented as the average ratio of pS6 or S6 to Tubulin ± SEM and are expressed as the percentage of water control. Significance was determined using one-way ANOVA. *n*_J=_J5-6, per condition. ns = non-significant.

We found that unlike male mice in which the basal levels of mTORC1 activity are consistently very low (Neasta, Ben Hamida et al. 2010, Laguesse, Morisot et al. 2017, Ehinger, Zhang et al. 2021), S6 phosphorylation and thus mTORC1 activation in water only drinking female mice is very variable, in some mice S6 phosphorylation was very low in several mice and very high in others (**Fig. 1c-d**). Interestingly, in contrast to mTORC1 being robustly activated by alcohol in the NAc of male mice both during 4 hours of alcohol binge drinking session and after 24 hours of alcohol withdrawal (Neasta, Ben Hamida et al. 2010, Laguesse, Morisot et al. 2017, Ehinger, Zhang et al. 2021), alcohol intake of female mice produced variable levels of S6 phosphorylation in the NAc (**Fig. 1c-d**). Specifically, binge drinking of alcohol produced a consistent reduction of S6 phosphorylation in the NAc of female mice, however the change was not significant due to the variability in S6 phosphorylation in the NAc of water consuming female mice (**Fig. 1c-d**). In contrast, S6 phosphorylation was elevated in 4 out of the 6 female mice at the end of the withdrawal period (**Fig. 1c-d**). Finally, similar to what was observed in male mice (Neasta, Ben Hamida et al. 2010, Laguesse, Morisot et al. 2017, Ehinger, Zhang et al. 2021), the total levels of S6 in the NAc were similar in water and alcohol consuming female mice (**Fig. 1c-d**).

We previously showed that heavy alcohol use produces a significant activation of mTORC1 in the OFC of male mice and rats (Laguesse, Morisot et al. 2017). We further showed that mTORC1 in the OFC contributes to alcohol seeking and habit but not drinking *per se* (Morisot, Phamluong et al. 2019). Thus, we measured the level of S6 phosphorylation in the OFC of females that have been consuming 20% alcohol for 7 weeks (**Fig. 1a**). The OFC was dissected after the last 4-hour binge drinking session and at the end of the last 24-hour withdrawal session (**Fig. 2a**). In contrast to the profile of mTORC1 activity in the NAc of female mice, the baseline of S6 phosphorylation in the OFC of water only consuming mice was less variable (**Fig. 2b-c**). Binge drinking did not alter S6 phosphorylation in the OFC of female mice (**Fig. 2b-c**). However, alcohol withdrawal produced a reduction of mTORC1 activation which was not significant due to the variability of S6 phosphorylation in the in the water only drinking female mice (**Fig. 2b-c**). Like in males, the level of total S6 was similar in female mice consuming water only or alcohol (**Fig. 2b-c**).

**Figure 2:**
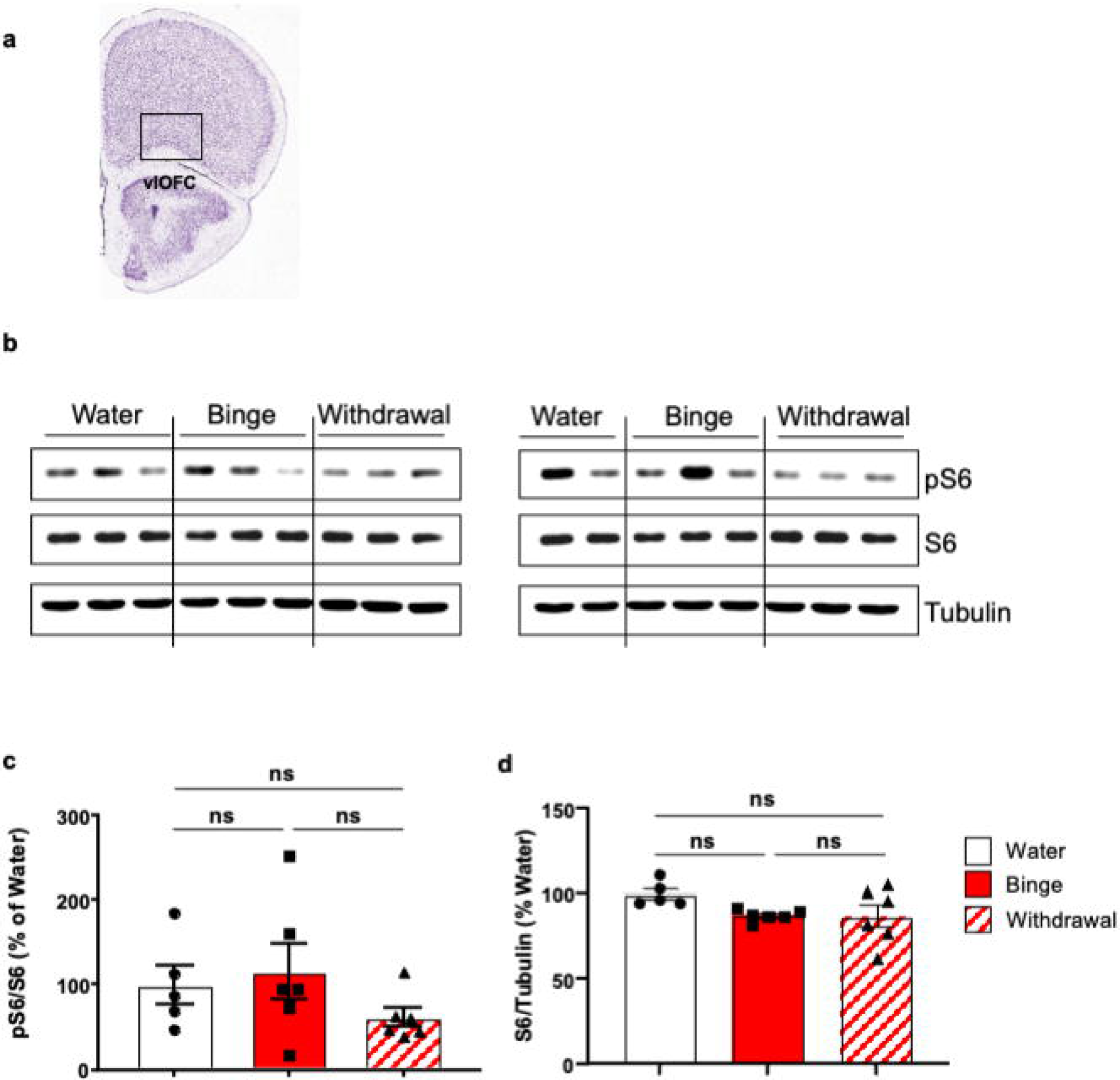
mTORC1 is not activated in the OFC of female mice in response to excessive alcohol intake. Mice underwent 7 weeks of IA20%-2BC whereas control animals had access to water bottles only. The orbitofrontal cortex (OFC) (**a**) was dissected after the last 4h binge drinking session or at the end of the last withdrawal period and was subjected to western blot analysis. (**c-e**) PhosphoS6 and S6 protein levels were determined by western blot analysis. Tubulin was used as a loading control. ImageJ was used for optical density quantification. Data are presented as the average ratio of pS6 or S6 to Tubulin ± SEM and are expressed as the percentage of water control. Significance was determined using one-way ANOVA. *n* _J=_J 5-6, per condition. ns = non-significant.

We next examined the pattern of mTORC1 activation in the mPFC. We previously showed that mTORC1 is not activated by alcohol drinking in the mPFC of male mice and rats (Laguesse, Morisot et al. 2017). Thus, we determined whether the same will be true in the mPFC of female mice consuming high levels of alcohol. The mPFC of female mice consuming water or 20% alcohol for 7 weeks was dissected during binge and withdrawal, and S6 phosphorylation was evaluated. Similar to the NAc of females drinking water only, the level of S6 phosphorylation in the mPFC of mice consuming water was variable; in some mice, a high level of S6 phosphorylation was detected. Binge drinking of alcohol and alcohol withdrawal produced a non-significant trend toward a reduction of mTORC1 activity in the mPFC of female mice (**Figure 3b-c**). The level of S6 was similar in alcohol drinking mice and was unchanged by alcohol (**Figure 3b-c**). Together, these data suggest that the pattern of mTORC1 activity in the 3 corticostriatal regions of female mice is vastly different from males during basal conditions in response to and during binge and withdrawal (data herein vs. (Neasta, Ben Hamida et al. 2010, Laguesse, Morisot et al. 2017, Ehinger, Zhang et al. 2021)).

**Figure 3:**
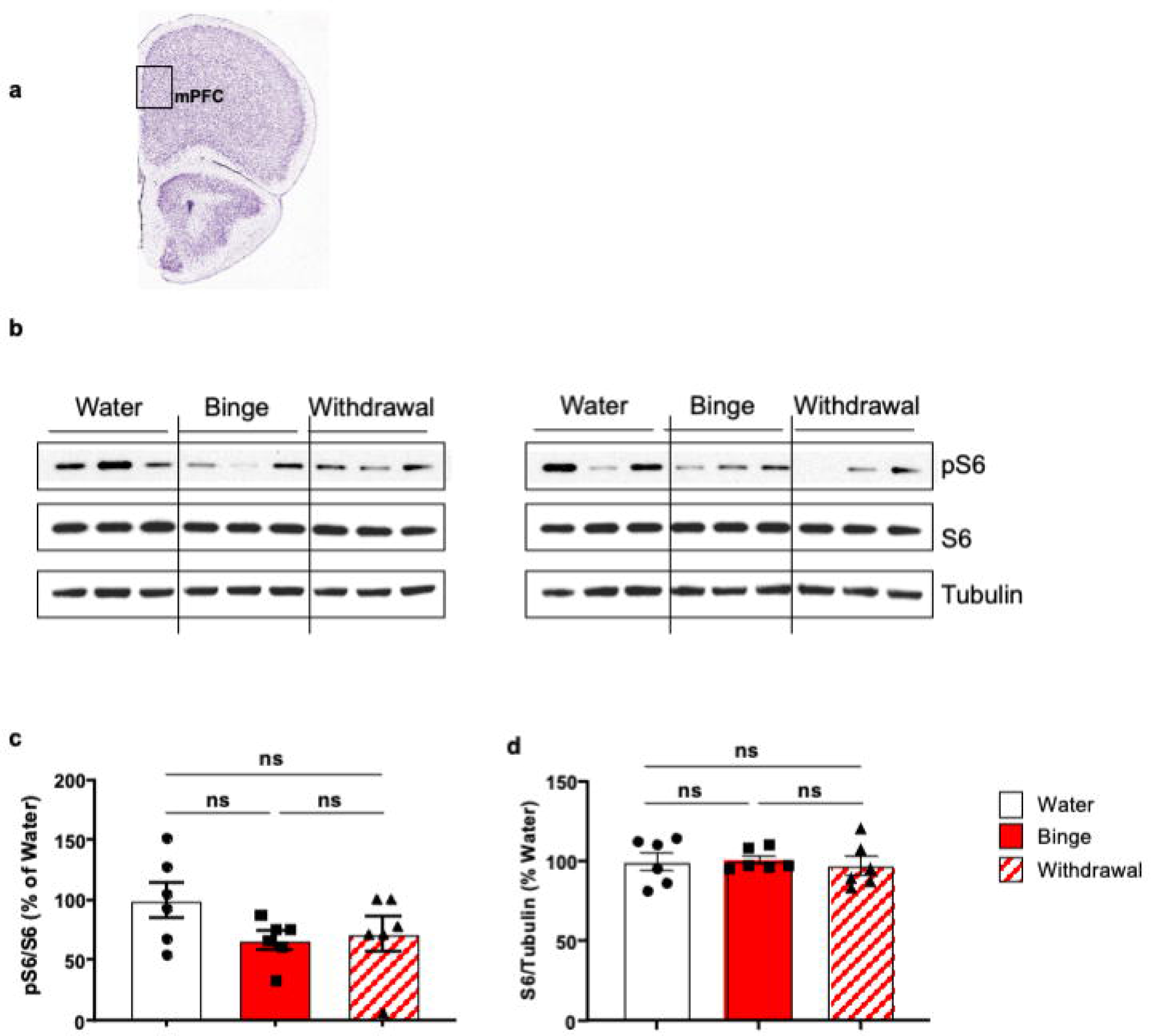
mTORC1 is not activated in the mPFC of female mice in response to excessive alcohol intake. Mice underwent 7 weeks of IA20%-2BC whereas control animals had access to water bottles only. The medial prefrontal cortex (mPFC) (**a**) was dissected either after the last 4h binge drinking session or at the end of the last withdrawal period and was subjected to western blot analysis. (**c-e**) PhosphoS6 and S6 protein levels were determined by western blot analysis. Tubulin was used as a loading control. ImageJ was used for optical density quantification. Data are presented as the average ratio of pS6 or S6 to Tubulin ± SEM and are expressed as the percentage of water control. Significance was determined using one-way ANOVA. *n*_J =_J 5-6, per condition. ns = non-significant.

We next investigated whether the level of phosphorylation across the 3 brain regions within the different conditions (water, binge, and withdrawal) were correlated. As shown in **Figure 4a**, in the basal level of S6 phosphorylation in the NAc and the OFC or mPFC did not correlate. However, we found a positive correlation between the OFC and the mPFC of water drinking female mice (**Fig. 4b**). We then focused on binge-drinking mice and found no correlation in the level of S6 phosphorylation between the NAc, OFC and mPFC s (**Fig. 4c**). Interestingly, we identified a positive correlation between the level of phosphor S6 in the mPFC and the average of the last week of drinking, suggesting that the drinking profile can predict the mTORC1 activation in the mPFC after binge drinking (**Fig. 4d**). Lastly, the level of S6 phosphorylation in the NAc, OFC and mPFC during withdrawal were not correlated (**Fig. 4e**).

**Figure 4:**
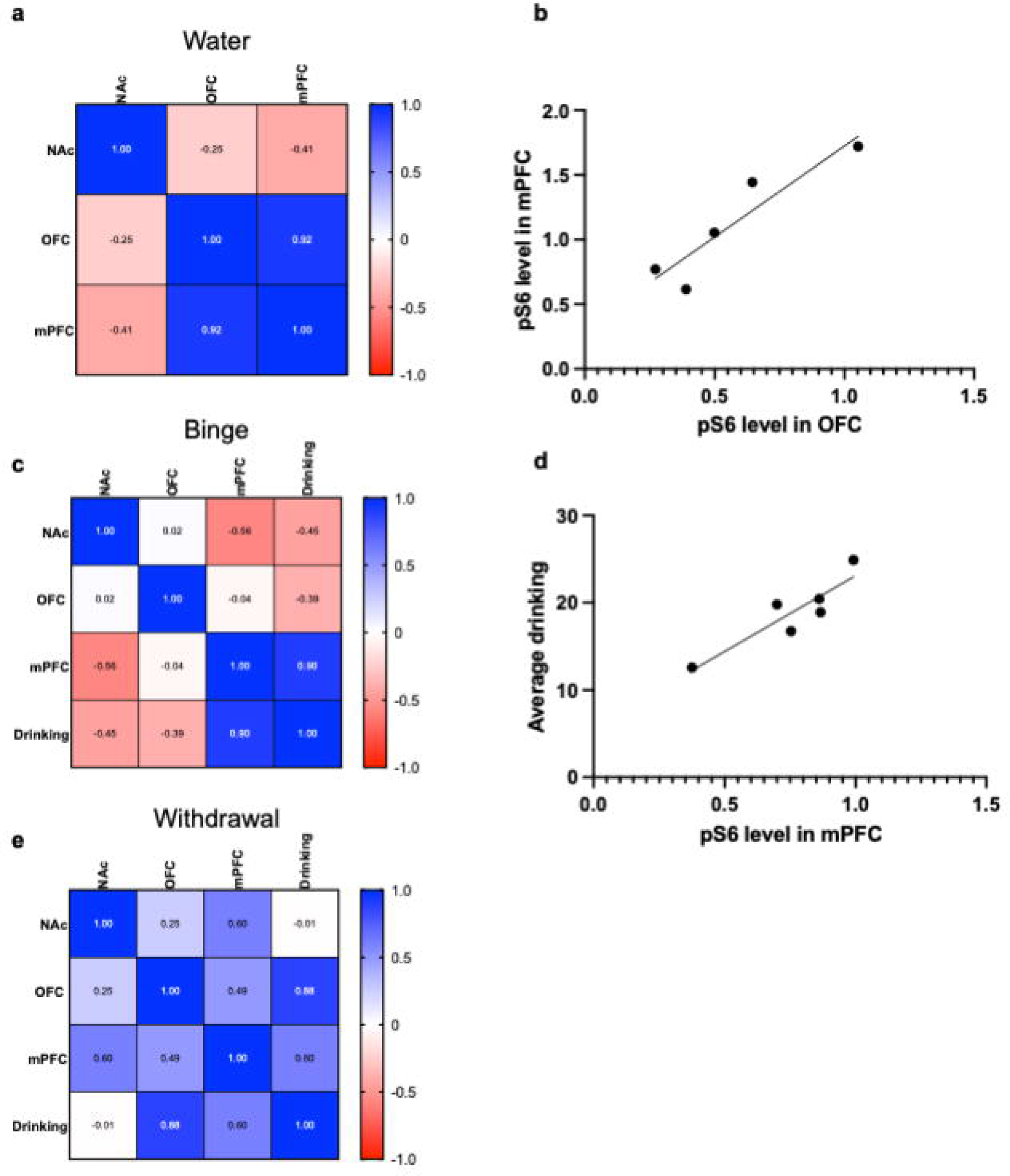
Pearson correlations comparison between brain region and conditions. (**a**) Heatmap summarizing Pearson’s correlation coefficients between the NAc, OFC and mPFC of water drinking mice. (**b**) Average alcohol drinking during the last week is positively correlated to the level of phosphoS6 in the mPFC after binge drinking (R2 = 0.8549, p = 0.0246). (**c**) Heatmap summarizing Pearson’s correlation coefficients between the NAc, OFC and mPFC after binge drinking. (**d**) level of phosphoS6 in the mPFC is positively correlated with the level of phosphoS6 in the OFC after binge drinking (R2 = 0.8165 p = 0.0135). (**e**) Heatmap summarizing Pearson’s correlation coefficients between the NAc, OFC and mPFC after 24h withdrawal. Positive correlations are indicated in shades of blue, whereas negative correlations are indicated in shades of red. *n* _J=_J 5-6, per condition. ns = non-significant.

Finally, we determined whether systemic administration of the selective mTORC1 inhibitor, rapamycin (Li, Kim et al. 2014), alters alcohol intake in female mice. To do so, female mice underwent 7 weeks of IA20%2BC reaching an average of 18.7 ± 1.41 g/kg/24hr of alcohol intake (**Table 1**). Rapamycin (20g/kg) was administered systemically 3 hours before the beginning of the drinking session (**Fig. 5a**). In contrast to male mice (Neasta, Ben Hamida et al. 2010, Morisot, Novotny et al. 2018, Ehinger, Zhang et al. 2021), systemic inhibition of mTORC1 did not alter females’ 4-hour or 24 hours drinking and preference (**Fig. 5b-e**). Water intake was also unaltered by rapamycin (**Fig. 5f-g**). Together, these data suggest that the interaction between alcohol and mTORC1 is sex-dependent, and that mTORC1 does not contribute to the neuroadaptations that underlie excessive alcohol intake in female mice.

**Figure 5:**
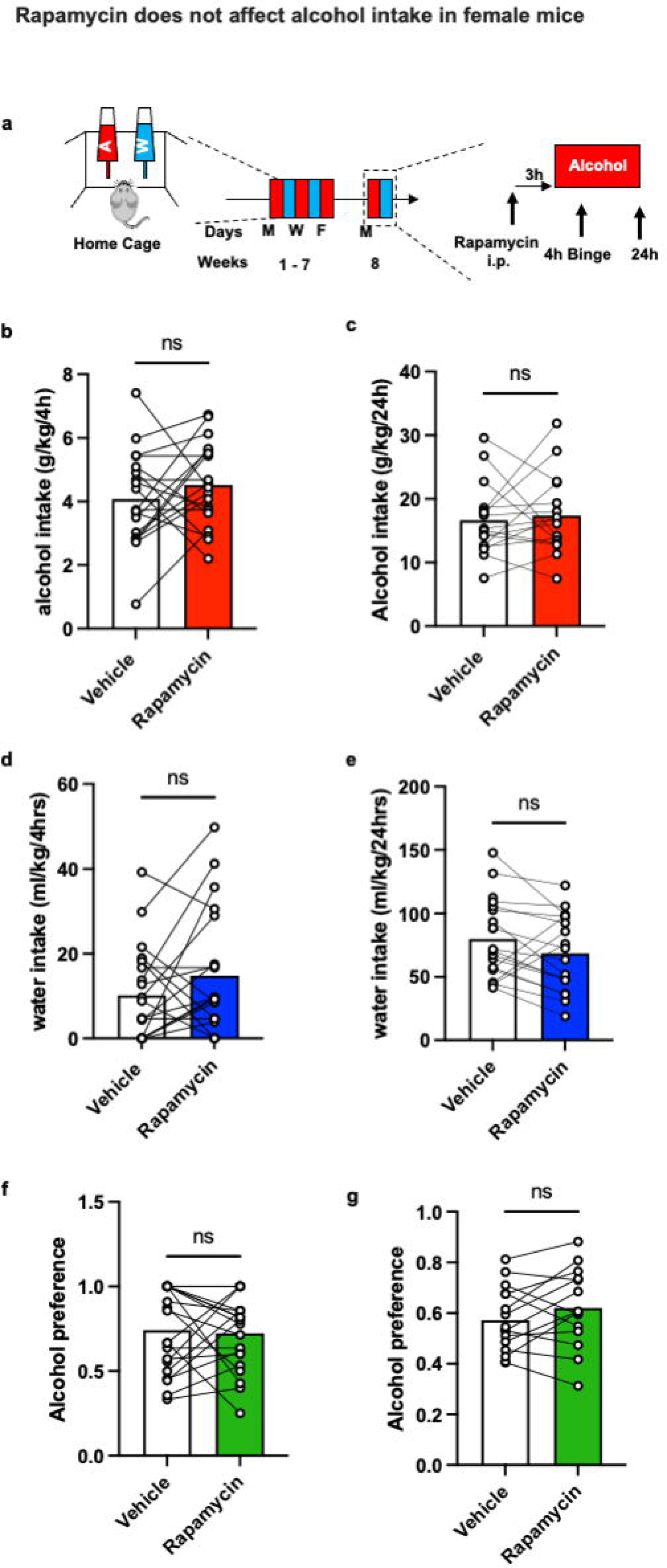
Rapamycin does not affect alcohol intake in female mice. **(a)** Timeline of experiment. Mice underwent 7 weeks of IA20%2BC. On week 8, mice received a systemic administration of vehicle (white) or Rapamycin (20_Jmg/kg, red) 3-hour before the beginning of drinking session. Alcohol and water intake were measured 4-hour (**b**, **d**) and 24_J hours later (**c**, **e**). Alcohol preference was calculated as the ratio of alcohol intake to total fluid intake at the end of the 4-hour (**f**) and 24_J-hour (**g**) drinking session. Significance was determined using a paired t-test. n _J=_J 20 per condition. ns = non-significant.

## DISCUSSION

Here, we provide evidence to suggest that intermittent access to 20% alcohol does not activate mTORC1 in the NAc, OFC and mPFC of female mice. We further show that systemic administration of rapamycin does not alter excessive alcohol intake and preference in females. These results are vastly different from what was obtained in male animals in which mTORC1 was robustly activated by alcohol in the NAc and OFC of male mice during binge drinking session and after 24 of withdrawal from alcohol intake (Neasta, Ben Hamida et al. 2010, Laguesse, Morisot et al. 2017, Ehinger, Zhang et al. 2021).

While evaluating mTORC1 activation state in females, several observations are noted. We detected a mouse-to-mouse variability in the basal levels of S6 phosphorylation and thus mTORC1 activity in the three brain regions. Another curious observation is that there was no correlation between a specific female mouse and its basal levels of S6 phosphorylation in the NAc when compared to the prefrontal brain regions. For example, we detected a low basal level of S6 phosphorylation in the NAc of a mouse drinking water and high level of S6 phosphorylation in the mPFC of the same mouse. In contrast, we identify a positive correlation in the basal level of phospho S6 in the OFC and mPFC, suggesting a potential common mechanism in mTORC1 regulation in the prefrontal cortex of female mice. We did not detect differences in the total levels of S6 within the individual female mice drinking water, as we previously reported in male mice (Neasta, Ben Hamida et al. 2010, Laguesse, Morisot et al. 2017, Ehinger, Zhang et al. 2021). Thus, the animal-to-animal variation in basal S6 phosphorylation cannot be due to differences in the basal level of the protein. Finally, our results also indicate that the basal level of mTORC1 activation state in the majority of female mice is much higher than in male mice (this study vs. (Neasta, Ben Hamida et al. 2010, Laguesse, Morisot et al. 2017, Ehinger, Zhang et al. 2021)). This observation is very curious as mTORC1, like most other kinases, is quiescent until an appropriate signal activates it. What could be the reason that the basal mTORC1 activity is high in female as compared to male mice? First, as our animals are socially isolated it is plausible that stress and/or social isolation activate mTORC1 in corticostriatal regions. Another possibility is that the estrus cycle determines the activation state of mTORC1. This conclusion stems from the finding that estradiol was reported to activate mTORC1 (Miller, Balko et al. 2011, Alayev, Salamon et al. 2016). Thus, it is plausible that the high levels in mTORC1 activity in some of the female mice is due to the estrus cycle. Therefore, the activation patterns of mTORC1 in the corticostriatal regions in water drinking female mice during the estrus cycle needs to be further explored.

In male mice we found that mTORC1 is activated in the NAc and OFC but not in other striatal or cortical regions including the mPFC in response to long-term heavy alcohol intake in male rats and mice (Laguesse, Morisot et al. 2017). In contrast, we detected a trend albeit non-significant towards a reduction in the level of active mTORC1 during binge (NAc), withdrawal (OFC), and both binge and withdrawal (mPFC). These findings are interesting as it is the opposite of what was observed in male mice in which activation of mTORC1 was detected in the NAc and OFC of alcohol drinking mice. Additional animals are required to determine if this reduction is statistically significant, and if so, this finding could, at least in part, explain why rapamycin does not alter alcohol intake in female mice.

What could be the underlying cause of the sex-different effects of alcohol on mTORC1 signaling? It is plausible that alcohol alters the signaling cascade upstream of mTORC1 differently in male vs. female mice. For instance, mTORC1 is activated by the H-Ras/PI3K/AKT signaling (Hay and Sonenberg 2004). We previously showed that alcohol activates H-Ras (Ben Hamida, Neasta et al. 2012), PI3K and AKT (Neasta, Ben Hamida et al. 2011) and mTORC1 (Neasta, Ben Hamida et al. 2010, Laguesse, Morisot et al. 2017, Ehinger, Zhang et al. 2021) in the NAc of male mice. It is plausible that one of the components in the H-Ras/PI3K/AKT signaling is not activated, and/or not properly localized in the tested regions of female mice thereby leading to the observed sex difference. Furthermore, the level of the phosphatase PTEN (phosphatase and tensin homolog deleted on chromosome 10) is significantly higher in the cortex of female vs. male mice (de Mello, Andreotti et al. 2020). PTEN is an endogenous inhibitor of PI3K and AKT (Parsons 2020). Thus, it is plausible that alcohol affects PTEN which in turn results in different patterns of alcohol-dependent activation or inhibition of PI3K/AKT/mTORC1 in alcohol drinking mice.

Sex chromosome genes have garnered attention in recent years for their potential roles in driving sex-specific differences in behavior and susceptibility to various conditions. Research involving mouse models with differing sex chromosome complements (XX, XY) and gonadal sex such as the four Core Genotypes mice has provided valuable insights (Corre, Friedel et al. 2016, Arnold 2020, Ghosh, Chen et al. 2021, Arnold, Chen et al. 2023, Cioffi, Grassi et al. 2024). These models allowed scientists to differentiate the impacts of sex chromosomes themselves from the influence of gonadal hormones (Corre, Friedel et al. 2016, Ghosh, Chen et al. 2021, Arnold, Chen et al. 2023, Cioffi, Grassi et al. 2024). Using these models, Sneddon et al. showed in a recent study that chromosomal sex plays a role in shaping the development of binge-like alcohol consumption and aversion-resistance drinking in females (Sneddon, Rasizer et al. 2022, Sneddon, Masters et al. 2023).

We found that systemic administration of rapamycin reduces alcohol intake in male mice (Neasta, Ben Hamida et al. 2010) and we show herein that rapamycin does not alter alcohol consumption of female mice. Cozzoli et al. previously reported that intra-NAc administration of rapamycin decreased binge drinking in male but not female mice (Cozzoli, Kaufman et al. 2016), confirming our own findings (Neasta, Ben Hamida et al. 2010). However, Cozzoli et al. study should be taken with a grain of salt. Specifically, the authors reported that the protein levels upstream and downstream of mTORC1 including PI3K, S6K, and 4EBP were reduced in the NAc response to binge drinking of male but not female mice (Cozzoli, Kaufman et al. 2016). These data are vastly different from ours in which we found no difference in the level of PI3K (Neasta, Ben Hamida et al. 2011), or S6K and 4EBP (Neasta, Ben Hamida et al. 2010). Furthermore, it is unclear how an inhibitor can reduce alcohol consumption when the signaling cascade is already attenuated by alcohol.

The fact that rapamycin does not inhibit alcohol drinking in female mice does not exclude the possibility that the kinase promotes other alcohol-dependent behaviors through its action in other brain regions associated with AUD such as the central amygdala (CeA), the basolateral amygdala (BLA), the hippocampus. It is also plausible that alcohol cues but not intake activates mTORC1 in female mice. For example, we previously found that reconsolidation of alcohol cue reward memories robustly activates mTORC1 signaling in the CeA and mPFC of male rats and that inhibition of mTORC1 via intra-CeA or systemic administration of rapamycin inhibits alcohol cue reward memories (Barak, Liu et al. 2013). Thus, it is plausible that mTORC1 signaling is activated in other brain regions in female mice which in turn promotes maladaptive phenotypes such as relapse.

A great deal is known by now about the differences in AUD onset, propensity, severity and associated phenotypes in males vs. females (Flores-Bonilla and Richardson 2020). Furthermore, a lot is known about how alcohol alters the molecular and cellular landscape in the brain of male rodents (Ron and Barak 2016, Egervari, Siciliano et al. 2021), however, much less is known about how alcohol alters molecular signaling in females. Our results highlight the importance of conducting such research. Furthermore, results such as ours are of importance as they indicate that one drug (e.g. rapamycin) does not fit all and point to the importance of personalized medicine.

## Acknowledgments

This research was supported by the National Institute of Alcohol Abuse and Alcoholism, R01 AA027474.

